# Triphasic production of IFNγ by innate and adaptive lymphocytes following influenza A virus infection

**DOI:** 10.1101/2023.05.23.541923

**Authors:** George E Finney, Kerrie E Hargrave, Marieke Pingen, Thomas Purnell, David Todd, Freya MacDonald, Julie C Worrell, Megan KL MacLeod

**Affiliations:** Centre for Immunobiology, School of Infection and Immunity, University of Glasgow, Glasgow, UK; Institute of Immunity and Transplantation, University College London, London, UK; Strathclyde Institute of Pharmacy and Biomedical Sciences, University of Strathclyde, UK

**Keywords:** immune memory, interferon-gamma, influenza A virus, lung, trained immunity

## Abstract

Interferon gamma (IFN*γ*) is a potent antiviral cytokine that can be produced by many innate and adaptive immune cells during infection. Currently, our understanding of which cells produce IFN*γ* and where they are located at different stages of an infection are limited. We have used reporter mice to investigate *in vivo* expression of IFN*γ* in the lung and secondary lymphoid organs during and following influenza A virus (IAV) infection. We observed a triphasic production of IFN*γ* expression. Unconventional T cells and innate lymphoid cells, particularly NK cells, were the dominant producers of early IFN*γ*, while CD4 and CD8 T cells were the main producers by day 10 post-infection. Following viral clearance, some memory CD4 and CD8 T cells continued to produce IFN*γ* in the lungs and draining lymph node. Interestingly, IFN*γ* production by lymph node Natural Killer (NK), NKT and innate lymphoid 1 cells also continued to be above naïve levels, suggesting memory-like phenotypes for these cells. Analysis of the localisation of IFN*γ*+ memory CD4 and CD8 T cells demonstrated that cytokine+ T cells were located near airways and in the lung parenchyma. Following a second IAV challenge, lung IAV specific CD8 T cells rapidly increased their expression of IFN*γ* while CD4 T cells in the draining lymph node increased their IFN*γ* response. Together, these data suggest that IFN*γ* production fluctuates based on cellular source and location, both of which could impact subsequent immune responses.

## Introduction

Interferon gamma (IFN*γ*) is a key cytokine that plays multiple roles in the host immune response to viral infections, for example influenza A virus (IAV). IFN*γ* promotes innate and adaptive leukocyte recruitment to the site of infection through the induction of CCR2 and CXCR3 ligands^1, 2^. Moreover, IFN*γ* signalling upregulates antigen presentation through major histocompatibility class I (MHCI) and MHCII pathways to CD8 and CD4 T cell respectively^3,4,5^ promoting T cell activation and ultimately facilitating viral clearance.

Various different immune cells have been documented to produce IFN*γ* following IAV infection. These include innate lymphoid cells (Natural Killer (NK) cells^6^, Innate Lymphoid 1 cells (ILC1s^7^)), unconventional T cells (NKT^8^ and *γδ* T cells^9, 10^) as well as classical *αβ* T cells (CD4 and CD8 T cells^11, 12^). However, the dynamics of when and how much IFN*γ* these cells can produce at different stages of IAV infection, and after re-infection, have not been comprehensively and simultaneously analysed.

IFN*γ* production by CD4 and CD8 T cells is correlated with protection from IAV infection in humans^13,14,15,16^. Most of these studies examine peripheral blood mononuclear cells, although, IFN*γ*+ CD8 T cells increase in the bronchial alveolar lavage fluid during challenge infections^17^. Similar findings have been observed in mouse models of IAV infection demonstrating that IFN*γ*+ CD4 and CD8 T cells can protect mice from challenge infection^18,19,20^. A major advantage of mouse infection models is the ability to examine immune cells within different organs at multiple time points following infection. This can provide a broader understanding of the cell types that are key IFN*γ* producers following IAV infection. A further advantage of mouse studies is the ability to identify cytokine producing cells via fluorescent reporter proteins^21, 22^. Thus, IFN*γ*+ cells can be detected without the need for *ex vivo* stimulation providing a more accurate view of the cells that respond to the virus *in vivo*.

Here, we characterised IFN*γ* expression at different timepoints following IAV infection within the spleen, mediastinal lymph node (Med LN), and in the lungs of IFN*γ* reporter mice. As expected, innate cells, particularly NK cells, were the most prominent IFN*γ* producers early post-infection, while conventional T cells were the largest IFN*γ* population at day 10. In contrast, following IAV re-challenge, T cells rapidly produced IFN*γ*, demonstrating their memory potential.

Interestingly, elevated IFN*γ* levels were sustained at day 40 post-infection, even though IAV is cleared by day 10^23, 24^. At this timepoint, CD4 and CD8 T cells were the only cells in the lungs that exhibited elevated IFN*γ* expression. In contrast, raised IFN*γ* expression at day 40 was found in ILC1s, NK and NKT cells, as well as conventional T cells, in the Med LN. The continued heightened production of IFN*γ* by non-adaptive immune cells echoes evidence that innate cells including ILC1s^25, 26^, NK^27^ and NKT cells^28^ can display memory-like properties and increased responsiveness to inflammatory stimuli.

Together, these data provide a comprehensive study of immune cell-derived IFN*γ* expression over the course of IAV infection. The importance of both local and peripheral IFN*γ* expression from different cell types may facilitate the design of interventions that either boost or inhibit cytokine production.

## Results

### Unconventional T cells and innate lymphoid cells produce IFN*γ* in the early and adaptive immune response to IAV infection in the lung and mediastinal lymph node

We used reporter mice, GREAT, to identify immune cells that express cytokines *in vivo* during and following an influenza A virus infection (IAV). These mice were crossed to IL-17 reporter mice, SMART, but in these experiments we have focussed on IFNγ producing cells. GREAT transgenic mice report IFNγ via live expression of EYFP^29^ which can be detected by flow cytometry (Gating shown in Supplementary Figure 1). We examined immune cells in the spleen, the lung draining, Med LN, and in the lung at early (day 5), peak adaptive immune response (day 10), and memory (day 40) timepoints^30^.

Following IAV infection, we could identify multiple populations of IFNγ+ immune cells (Supplementary Figure 2-3). By injecting the animals with fluorescently labelled anti-CD45 shortly before euthanasia, we could distinguish cytokine producing cells present in the blood from those in the tissues^31^. We have focussed on the cells in the lung tissue rather than blood, as these are the cells that are most likely to contributing to viral control and clearance.

The majority of EYFP+ cells were cells known to produce IFNγ from previous studies: NK cells, NKT cells, ILC1s, *γδ* T cells, CD4 and CD8 T cells^6,7,8,9,10,11,12^ (Supplementary Figure 4). Smaller populations were positive for our ‘dump’ antibodies which included B220, MHCII and F4/80, suggesting a small proportion of B cells and/or DCs and macrophages can produce IFNγ. A small percentage (1-2%) of IFN*γ*+ cells were not included in any of the gates shown in Supplementary Figure 1.

To track the dynamic changes in the main IFNγ+ populations during IAV infection, we first quantified the total numbers and numbers of IFNγ+ innate and unconventional T cells over time (Figure 1A-B, Supplementary Table 1-2s).

**Figure 1.**
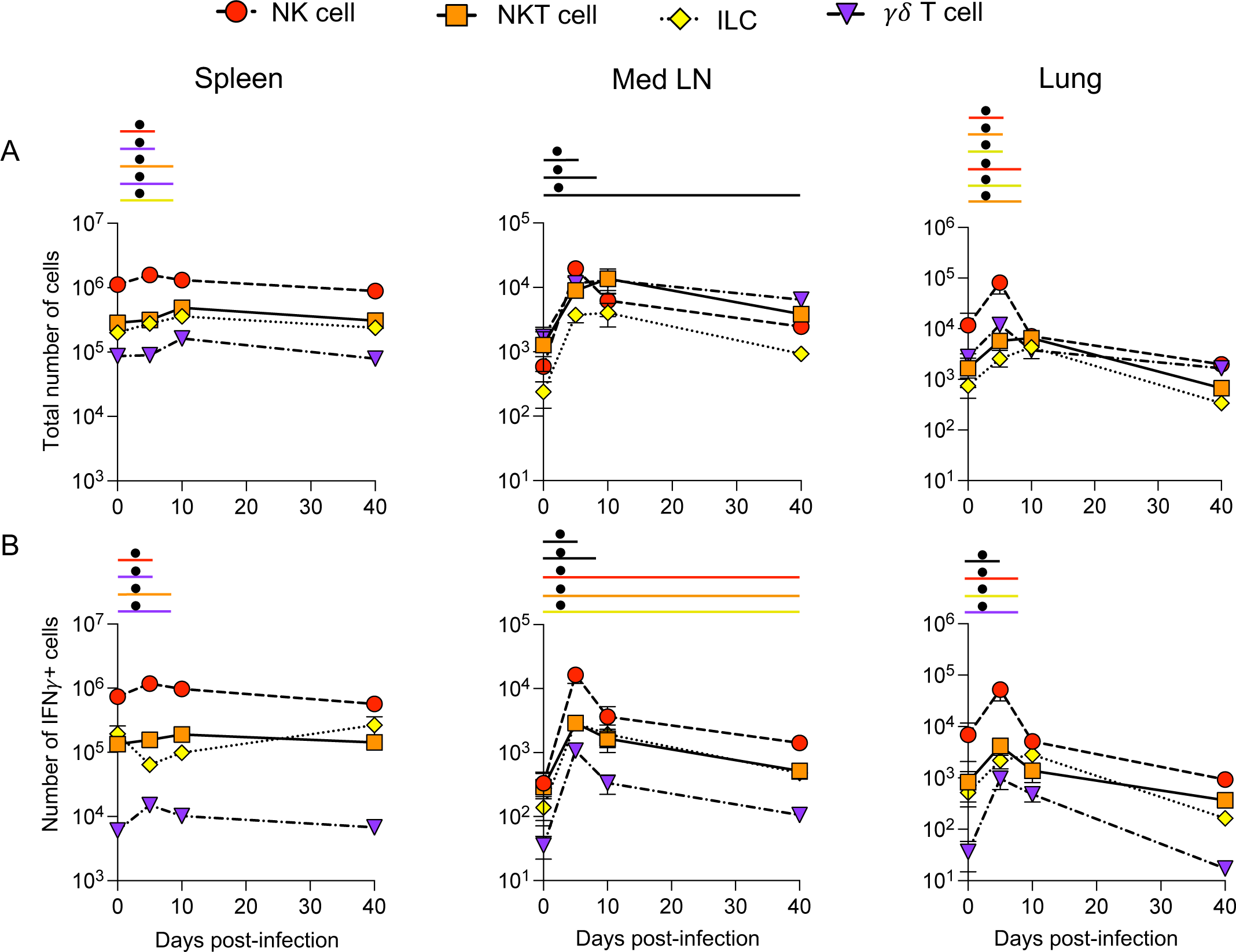
IFN*γ* production from unconventional T cells and innate lymphoid cells occurs early after IAV infection. GREATxSMART mice were infected with IAV on day 0 and injected with fluorescently labelled anti-CD45 i.v. 3 minutes prior to removal of the spleen, Med LN and lung. The total number (top, A) or number of IFN*γ*+ cells (bottom, B) of the indicated cell populations were examined in naïve animals or those infected 5, 10 or 40 days previously. Each point represents the mean of 8-11 infected mice from two independent time course experiments; 21 naïve mice are combined from across the time points and experiments, error bars are SEM. Coloured lines above each timepoints relates to significantly increased number of total cells or IFN*γ*+ cells compared to naïve animals. If line is black, this represents all cell types were higher at the indicated time point compared to naïve mice. Significance tested via a Kruskal–Wallis test followed by a Dunn’s multiple comparison test, •: significant difference between indicated timepoint and naïve samples, refer to table 1 & 2 for significance value.

There were minor, but in some cases significant, changes in the numbers of NK, NKT, ILCs and *γδ* T cells in the spleen following IAV infection (Figure 1A). The numbers of IFNγ+ NK, NKT and *γδ* T cells did increase significantly at either day 5 or day 10 post-infection but these changes were small (Figure 1B). At all timepoints, NK cells were the largest population of splenic IFNγ+ cells.

IAV infection led to more substantial changes in the total populations and numbers of IFN*γ*+ cells in the Med LN. Here, there were significant increases in the numbers of NK cells, NKT cells, ILC1s, and *γδ* T cells at all time points post-infection compared to naïve animals (Figure 1A). These results were also reflected in the numbers of IFNγ+ cells. IFN*γ*+ cells from all populations were increased at days 5 and 10, and all but *γδ* T cells remained increased at day 40 despite clearance of IAV by day 10^23, 24^ (Figure 1B). These data suggest a sustained immune response to IAV and evidence for innate or trained memory. As in the spleen, NK cells were the largest population of IFNγ+ cells within these cell types.

In the lung tissue, the numbers of ILC1, NK and NKT cells were increased 5 days after IAV infection compared to naïve animals, Figure 1A. While the numbers of ILC1s and NKT cells remained elevated in the lung 10 days post-infection, all four cell types returned to naïve levels by day 40 post-infection.

The numbers of IFNγ+ cells within each of the four populations were increased at day 5 post-infection and ILC1 and *γδ* T cells remained elevated at day 10. Notably the numbers of total NK cells and NKT cells dropped substantially by day 10 and were slightly slower than in naïve animals. In contrast to the sustained IFNγ in the Med LN, the number of IFNγ+ unconventional T cells and innate lymphoid cells returned to naïve levels by day 40 post-infection.

Together, these data suggest that of the innate cells, NK cells are the predominant source of IFNγ in the spleen, Med LN and lung. As expected, in the spleen and lung, the numbers of IFNγ+ cells returned to naïve levels of IFNγ by day 40 post-infection. In contrast, in the Med LN, we found evidence for long-term changes to several innate immune cell populations.

### CD4 and CD8 T cells are the predominant source of IFNγ after the clearance of IAV infection

T cells are key IFNγ producing populations during IAV infection^11, 12^. In the same experiments, therefore, we examined changes in IFNγ expression by CD4 and CD8 T cells over the course of IAV infection. There were minor changes in the numbers of total CD4 and CD8 T cells in the spleen after IAV infection, Figure 2A-B. While the numbers of IFNγ+ CD4 T cells remained stable, IAV infection did lead to a sustained increased in IFN*γ*+ CD8 T cells up to 40 days post-infection, Figure 2B.

**Figure 2.**
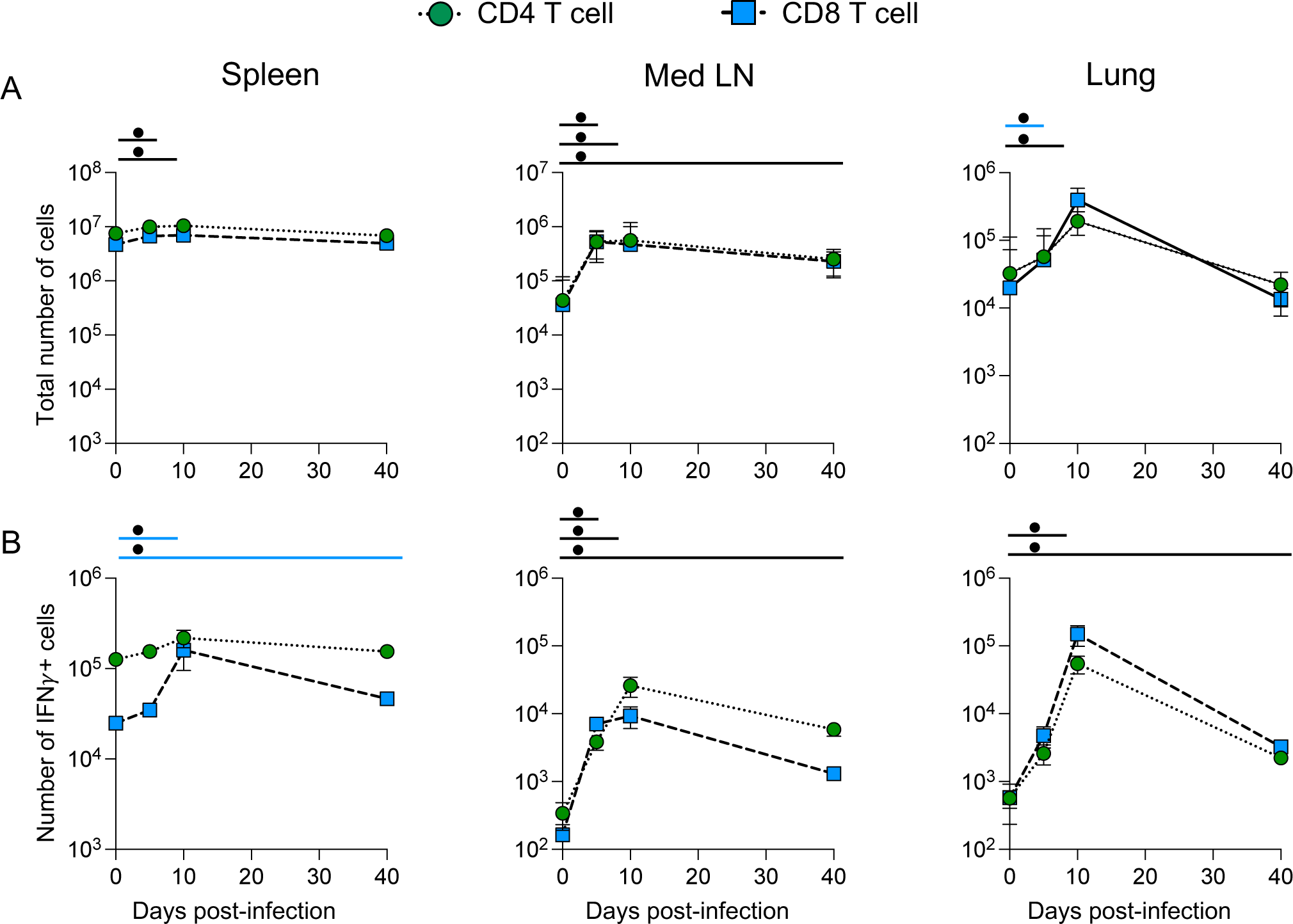
CD4 and CD8 T cells are the predominant source of IFN*γ* after the clearance of IAV infection. GREATxSMART mice were infected with IAV on day 0 and injected with fluorescently labelled anti-CD45 i.v. 3 minutes prior to removal of the spleen, Med LN and lung. The total number (top, A) or number of IFN*γ*+ cells (bottom, B) of the indicated cell populations were examined in naïve animals or those infected 5, 10 or 40 days previously. Each point represents the mean of 8-11 infected mice from two independent time course experiments; 21 naïve mice are combined from across the time points and experiments, error bars are SEM. Coloured lines above each timepoints relates to significantly increased number of total number or IFN*γ*+ cells compared to naïve animals of a specific cell type. If line is black, this represents all cell types were higher at the indicated time point compared to naïve mice. Significance tested via a Kruskal–Wallis test followed by a Dunn’s multiple comparison test, •: significant difference between indicated timepoint and naïve samples, refer to table 1 & 2 for significance value.

In the Med LN, at all time points following infection, the numbers of total cells and numbers of IFNγ+, CD4 and CD8 T cells were increased compared to naïve animals (Figure 2A-B). The numbers of IFNγ+ T cells peaked at day 10 post-infection with simialr numbers of cytokine+ CD4 and CD8 T cells. At day 40, their numbers remained above those in naïve animals (Figure 2B).

Within the lung tissue, there were no increases in the numbers of IFNγ+ CD4 and CD8 T cells 5 days after infection. By day 10, the numbers of total T cells and IFNγ+ T cells increased substantially and then declined by day 40. In contrast to the innate and unconventional T cells, the numbers of IFNγ+ CD4 and CD8 T cells did remain above naïve levels at day 40, indicating persistence of effector cytokine producing memory T cells. Additionally, by comparing the numbers of IFNγ+ cells from the different populations in the Med LN and lung at days 10 and 40 post-infection, we found that CD4 and CD8 T cells were the largest population of cytokine+ cells (Supplementary Figure 5).

We also examined the mean fluorescence intensity (MFI) of the EYFP signal to determine the relative amounts of IFNγ produced by the different lymphocytes across the time course (Supplementary Figure 6). For most of the innate cell types, there were limited, but in some cases significant, changes in the EYFP MFI in the spleen. Interestingly, CD4 and CD8 T cells expressed low levels of IFNγ at day 10 in the spleen and Med LN, suggesting these cells may receive a negative feedback signal at this time or the high cytokine+ cells may have migrated to the lung. At day 5 post-infection, in parallel with the increase in number of IFNγ+ ILC1, NK and NKT cells in the Med LN and lung, these cells increased the amount of IFNγ they produced. Similarly, lung CD4 and CD8 T cells increased their expression of IFNγ at day 10 post-infection, matching their increase in cell number.

To confirm that the IFNγ reporter marked cells that still expressed IFNγ transcript at the memory timepoints, we FACS sorted EYFP+ and negative CD44high CD4 and CD8 T cells from the Med LN and lungs of IAV infected mice. The EYFP+ CD4 T cells from these organs expressed more *Ifnγ* transcript than non-EYFP+ cells (Supplementary Figure 7). In contrast, the CD8 EYFP+ cells expressed similar transcript levels to EYFP negative cells and the transcripts levels were lower than for the CD4 T cells. Potentially there may be a technical reason for this difference, including a loss of *Ifnγ* transcript *ex vivo* by the CD8 T cells. Alternatively, *Ifnγ* transcript may be more rapidly degraded by CD8 than CD4 T cells *in vivo*.

### EYFP+ CD4 and CD8 T cells are located near airways and in the lung parenchyma

We used immunofluorescence to investigate the location of the lung CD4 and CD8 T cells that continue to produce IFNγ. We hypothesised that these cells would be located near the airways, ready to respond to a subsequent infection. CD4 and CD8 T cells were located both near EpCAM+ airways and in the parenchyma and this varied between animals (Figure 3). These data suggest that during the maintenance of T cell memory, there is no specialised niche in which the IFNγ cytokine+ T cells are located.

**Figure 3.**
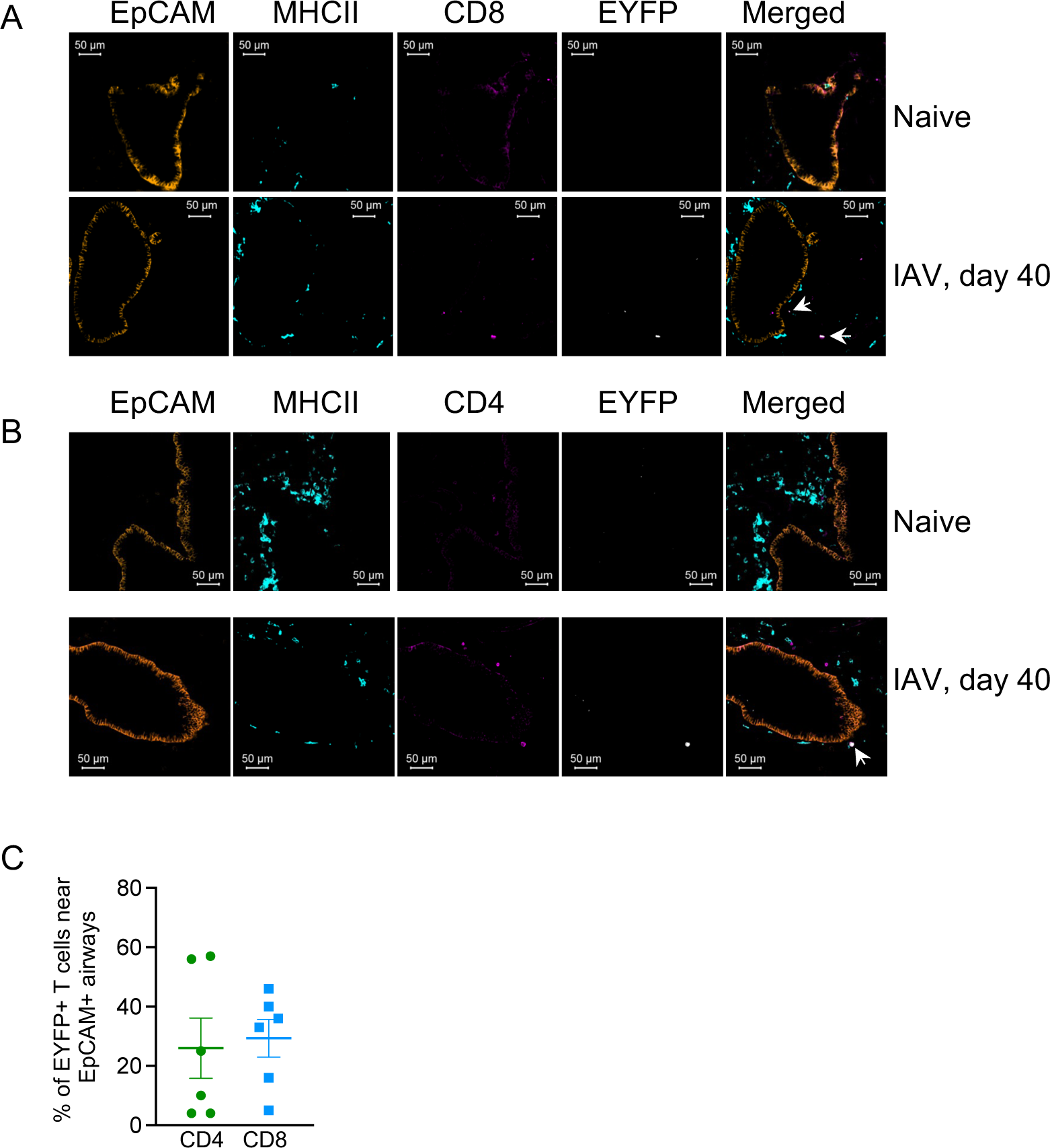
EYFP+ CD4 and CD8 T cells are located near airways and in the lung parenchyma. GREATxSMART mice were infected with IAV on day 0 and lungs analysed after 40 days. Example images of lung EYFP (white) CD8 (A) and CD4 (B) cells (pink) cells are shown alongside EpCAM+ cells (orange) and MHCII+ cells (turquoise) in naïve and infected animals and the percentages of CD4 or CD8 T cells either in close proximity to EpCAM+ airways or within the parenchyma determined in infected mice. In C, each symbol represents one mouse, data combined from 3-4 slides per animal. Data are combined from two experiments with 3 mice per experiment.

### IAV-specific CD4+ T cells continue to produce IFN*γ* following viral clearance

To confirm that IAV specific CD4 and CD8 T cells were producing IFNγ, we stained cells from IAV infected reporter mice with MHC I and MHC II tetramers containing immunodominant IAV nucleoprotein (NP) peptides, NP_368-74_ and NP_311-325_ respectively at days 10 and 40 post-infection (Figure 4A and 4B).

**Figure 4.**
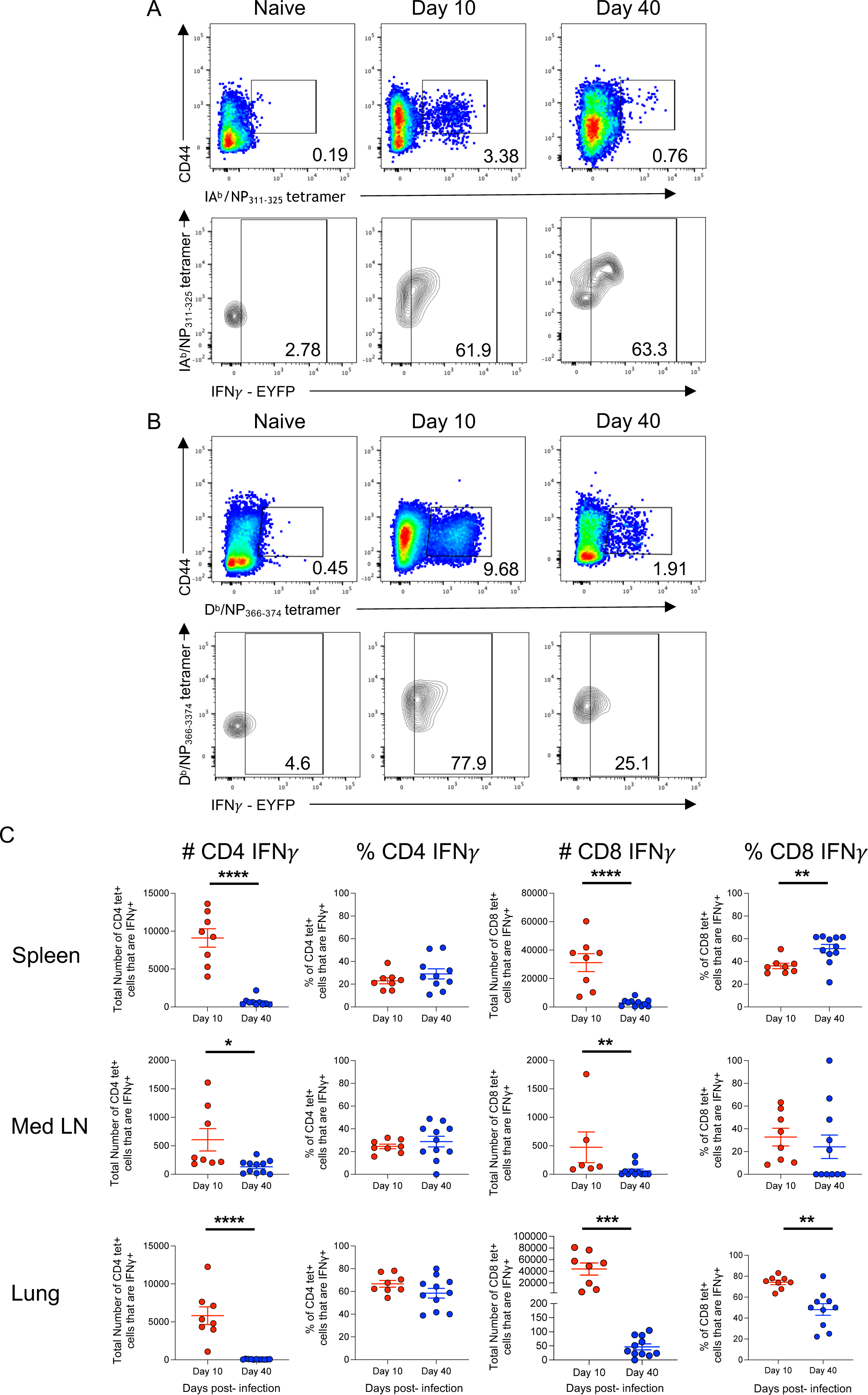
IAV-specific memory CD4+ T cells maintain the ability to transcribe IFN *γ* after viral clearance. GREATxSMART mice were infected with IAV on day 0 and injected with fluorescently labelled anti-CD45 i.v. 3 minutes prior to removal of the spleen, Med LN and lung. Representative flow plots display staining of (A) IAV-specific IFN*γ*+ CD4 or (B) CD8 T cells from the lung. (C) The frequency (left) and total number (right) of IAV-specific CD4 and CD8 T cells were compared between the peak T cell response (day 10; red) and after viral clearance (day 40; blue). Each point represents an individual mouse from 8-11 mice combined from two independent experiments; error bars are SEM. Significance tested via an unpaired t test, *: p<0.05, **: p<0.01, ***: p<0.001, ****: p<0.0001.

In the spleen, Med LN and lung, the numbers of IFNγ+ IA^b^/ NP_311-325_ tetramer+ CD4 and D^b^/NP_368-374_ tetramer+ CD8 T cells declined between the peak of the T cell response (day 10) and the memory time point (day 40) (Figure 4C). This decline in T cell number coincides with the vast majority of effector T cells dying by apoptosis, and the formation of long-lived memory T cells^32, 33^.

We also compared the percentages of CD4 and CD8 T cells that were IFNγ between day 10 and day 40 (Figure 4C). For IAV specific CD4 T cells, there was no differences in any of the organs. This suggests that cells that express IFNγ are as likely as non-IFNγ+ cells to enter the memory pool. Alternatively, memory CD4 T cells may fluctuate in their ability to express IFNγ depending on their location. In the spleen, 29% (±5.5%), and Med LN, 29% (±5.8%), of the MHCII tetramer+ cells were IFNγ+. In contrast, 59%(±3.5%) of the MHCII tetramer+ cells were IFNγ+ in the lung.

At day 10 post-infection, the majority of CD8 MHCI tetramer+ cells in the lung were IFNγ+. This dropped by approximately half by day 40 suggesting that non-IFNγ+ CD8 T cells are more likely to enter the memory pool. In contrast, in the spleen, there was a greater percentage of IFNγ+ CD8 MHCI tetramer+ cells at day 40 than day 10, while in the Med LN, there were no differences between day 10 and 40. These data may suggest that entry into the memory pool for CD8 T cells may follow different rules depending on the cell’s migration ability and/or reflect changes in memory cell function depending on location at the time of analysis.

### IFN*γ* lung NK cells and ILCs in the lung rapidly increase following re-infection and

We next examined the impact of a second IAV infection on IFN*γ* producing immune cells. Reporter mice were infected with WSN IAV (H1N1) and 30 days later some of these animals were re-infected with X31 (H3N2) and responding cells were analysed after a further 3 days.

We focussed on the innate and unconventional T cells in the lung as IFN*γ* is implicated in early IAV contro^18, 34^. In comparison to naïve animals, we only found increased numbers of IFN*γ*+ NK cells and ILCs in mice re-challenged with IAV, suggesting cytokine from these cells may be involved in early viral control (Figure 5).

**Figure 5.**
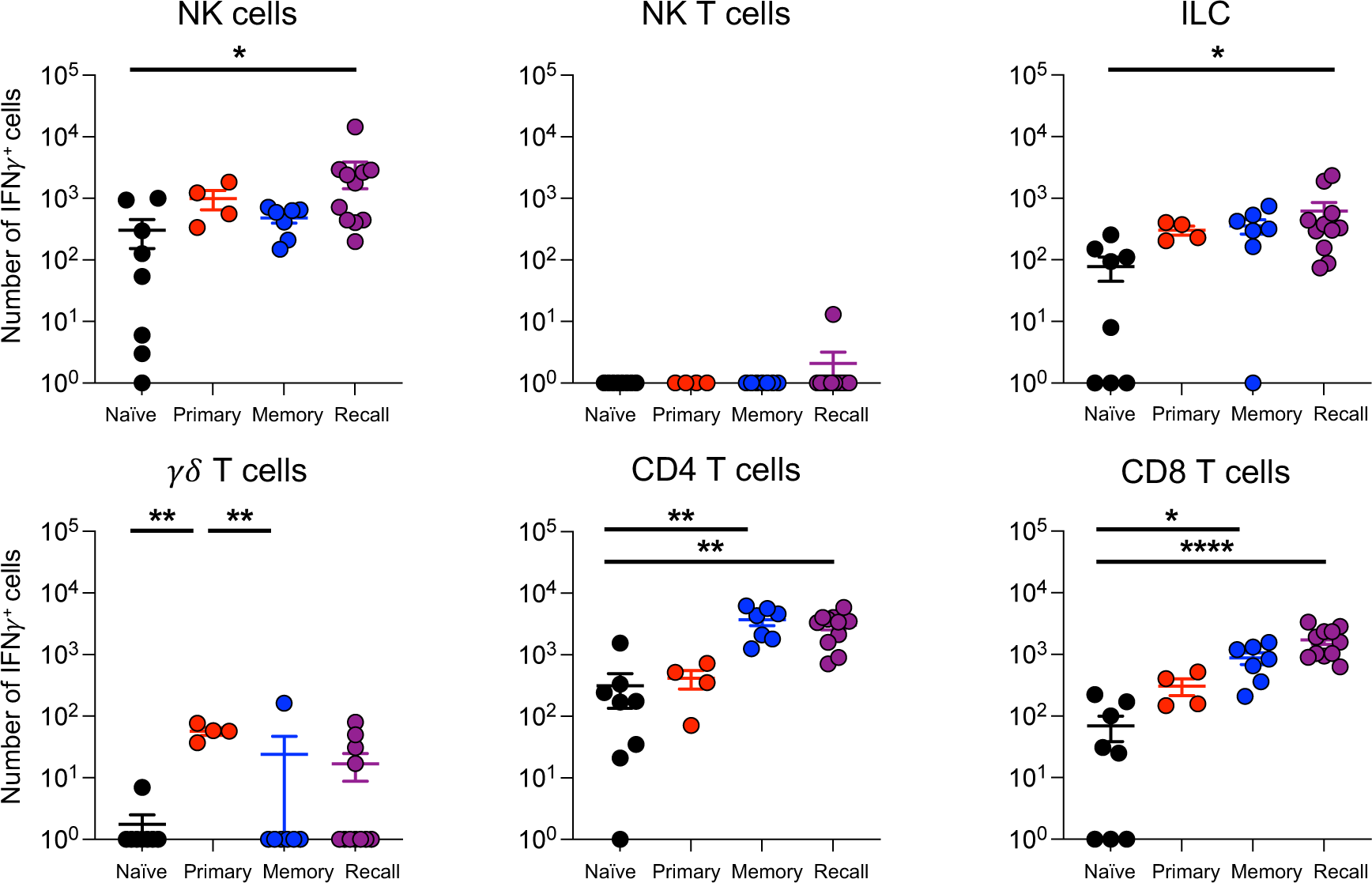
IFN*γ* lung NK cells and ILCs in the lung rapidly increase following re-infection. Naïve (primary) or WSN-infected GREATxSMART (recall) mice were infected i.n with X31 for 3 days or left unchallenged (naïve and memory). The numbers of IFN*γ*+ cells of immune cells in lung were examined by flow cytometry. Each point represents an individual mouse from 4-11 mice combined from two independent experiments; error bars are SEM. Significance tested via Kruskal-Wallis test followed by a Dunn’s multiple comparison, *: p<0.05, **: p<0.01, ****; p<0.0001.

### IAV-specific CD4 and CD8 T cells are more activated and increase in IFN*γ* expression following recall infection

While we did not observe changes in the numbers of IFN*γ*+ CD4 and CD8 T cells between memory and re-infected animals (Figure 5), by examining the antigen-specific T cell pools we found clear evidence for the re-activation of IAV specific T cells (Figure 6).

**Figure 6.**
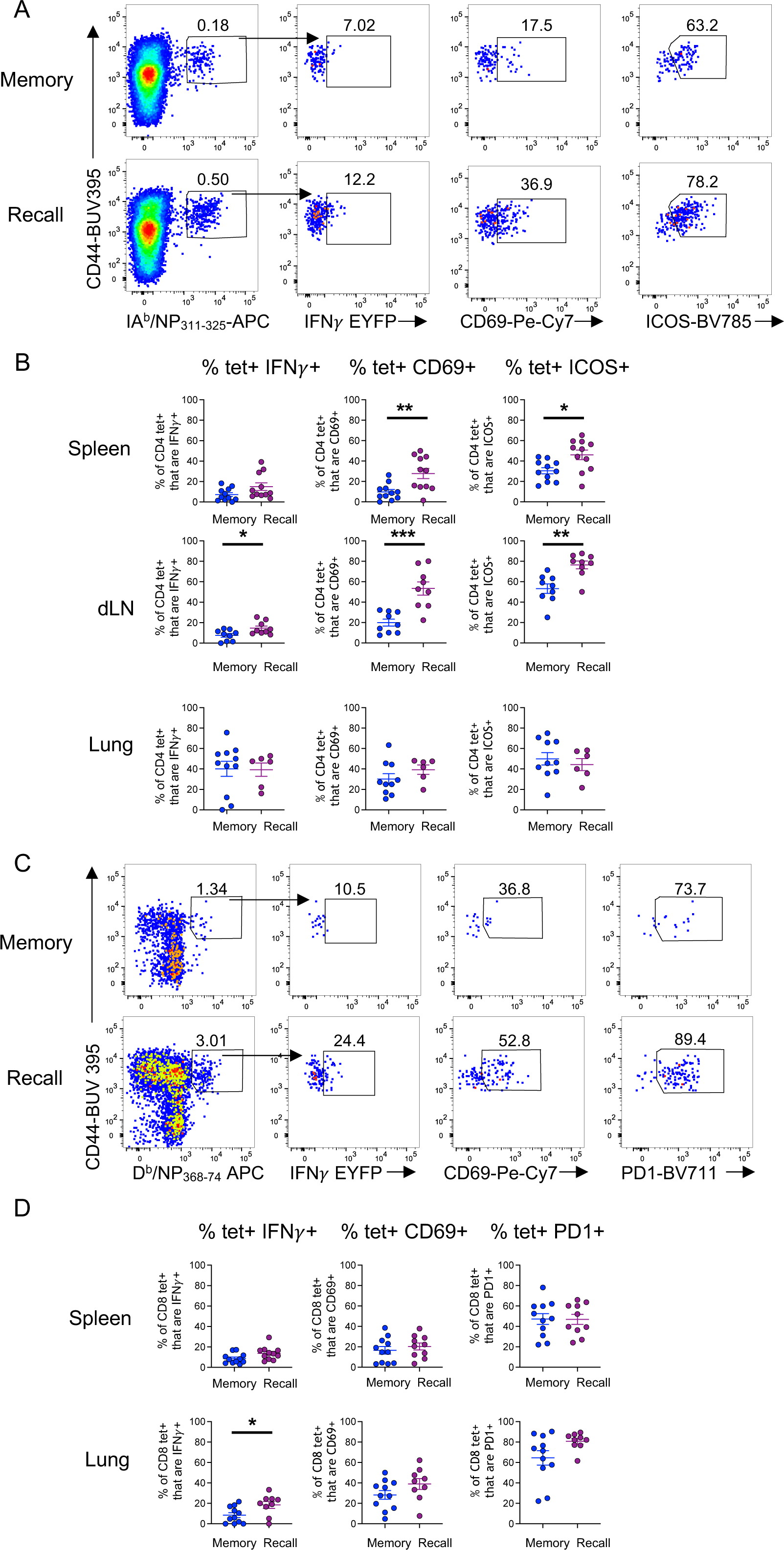
IAV-specific CD4 and CD8 T cells are activated and increase IFN*γ* expression following recall infection. GREATxSMART mice were either infected i.n with IAV WSN (H1N1) for 40 days (memory) or re-infected with X31 (H3N2) for 3 days. Antigen-specific CD4 and CD8 T cells from the lung, spleen and Med LN were examined. (A) Representative flow plots of IA^b^/NP_311-325_ specific CD4+ T cells from the Med LN in memory (top) and recall infection (bottom) and expression of (B) IFN*γ*, CD69 and ICOS from IA^b^/NP_311-325_ specific CD4+ T cells in the spleen, Med LN and lung. (C) Representative flow plots of D^b^/NP_368-374_ specific CD8+ T cells from the lung in memory (top) and recall infection (bottom) and expression of (D) IFN*γ*, CD69 and PD1 from D^b^/NP_368-374_specific CD8+ T cells in the spleen and lung of memory and recall. Each point represents an individual mouse from 9-11 mice combined from two independent experiments; error bars are SEM. Significance tested via an unpaired t test, *: p<0.05, **: p<0.01, ***: p<0.001. Some samples were removed from these groups for technical reasons: too few MHC tetramer+ cells, thymus contamination in the Med LN, or loss of cells during analysis.

After re-infection, Med LN IAV-specific CD4 T cells increased their expression of IFN*γ* and were more activated compared to IAV-specific CD4 T cells from memory mice based on increased expression of CD69 and ICOS (Figure 6A-B). Splenic IAV specific CD4 T cells were also activated by the re-infection, increasing their expression of CD69 and ICOS, although the expression of IFN*γ* was equivalent to that in memory animals.

IAV-specific CD4 T cells in the lung did not display increased expression of CD69, ICOS or IFN*γ* following re-infection, Figure 6B. However, we found that IAV specific lung CD4 T cells were more likely to express these molecules than those in spleen or Med LN prior to infection suggesting these cells remain in a semi-activated state following the initial infection.

We also compared the IFN*γ* expression and activation state of D^b^/NP_368-374_ tetramer+ CD8 T cells in the lung and spleen of memory and recall mice. There were no changes in splenic IAV specific CD8 T cells and too few tetramer+ CD8 T cells in the Med LN for analysis. There was, however, a significant increase in IFN*γ* expression from D^b^/NP_368-374_ tetramer+ CD8 T cells in the lungs following re-infection (Figure 6C-D). These cells showed some evidence for increased expression of CD69 and PD1, although these changes were not significant.

## Discussion

Our data demonstrate triphasic *in vivo* IFN*γ* production following IAV infection. Unconventional T cells and innate lymphoid cells were early cytokine producers, prior to the expansion and infiltration of IFN*γ* effector CD4 and CD8 T cells in secondary lymphoid organs and the lung. After viral clearance, IFN*γ* levels remained above naïve levels in a number of cell types, including adaptative and innate immune cells, for at least 40 days.

Our finding that NK cells provide early IFN*γ* production agrees with studies from Stegemann-Koniszewski and Wang^6, 35^. We have extended these data by demonstrating that NK cells are the dominant producers of IFN*γ* during early IAV infection, both in terms of numbers of cells and amount of cytokine produced. We also provide evidence that NKT cells, ILC1s and *γδ* T cells also produce IFN*γ* early during infection as other studies have described^7,8,9,10^. As expected, large numbers of CD4 and CD8 T cells produced IFN*γ* at day 10 post infection^30, 36^. Unexpectedly, we found that a number of cell types continued to express IFN*γ* at levels above naïve animals for at least 40 days. In the Med LN, these included CD4 and CD8 αβT cells, NKT cells, NK cells and ILC1. In all cases the total numbers of these cells were also higher at day 40 post-IAV than in naïve animals suggesting either continued recruitment to, or retention within, the lymph node.

While long-term alterations to conventional T cells following viral challenge are to be expected, classically, unconventional T cells and innate immune cells do not display memory characteristics. This view has been challenged in the last 10-20 years, in particular in the fields of NK cells and monocytes^37, 38^. In IAV infection studies, Li *et al* identified protective NK cells in the liver, but not the lung, of infected animals^43^. In contrast, Zheng *et a*l describe an alteration to splenic NK cells following IAV infection, leading to reduced cytotoxicity and an inability to protect adoptive immunodeficient hosts^44^. We found a rapid recruitment of cytokine+ cells to the lungs following a secondary infection. The response may be a result of the increased IFN*γ*+ NK cells in the lymph node after initial infection. Alternatively, the lung

CD4 and CD8 T cell response to re-infection could lead to enhanced migration of NK cells to the lung. Such collaboration between different types of immune cells would demonstrate a co-ordinated response that would be difficult to unpick using traditional single adoptive transfer approaches.

An alternative explanation for the continued elevated expression of IFN*γ* by multiple cell types is that persistent IAV antigens fuel a low level immune response. Studies from Jelly-Gibbs *et al* and Zammit *et al* demonstrate the presentation of IAV antigens to CD4 and CD8 T cells respectively for 1-2 months post infection^39, 40^. The continued presentation to conventional T cells could drive IFN*γ* expression which in turn supports the cytokine production by the unconventional and innate T cells, for example via promoting IL-12 production^41^.

Zammit *et al* tracked the persistent IAV antigen to the Med LN, but we found that T cells in the lung also continued to express IFN*γ* to day 40 post-infection. Antigen could also be retained in the lung, for example within clusters of immune cells^42, 43^. Various APC populations have been implicated in presenting antigen to tissue-resident memory cells including different dendritic cell populations^44, 45^, upper and lower airway epithelial cells^44, 45^, and lung fibroblasts^46^. From our imaging data, we found that IFN*γ*^+^ memory CD4 and CD8 T cells were located near EpCAM+ airways and in the parenchyma. This perhaps suggests that if persistent antigen drives this IFNγ response, a number of different cell types could drive this cytokine response.

In summary, our data demonstrate a triphasic production of *in vivo* IFN*γ* production, started by unconventional T cells and innate lymphoid cells early in infection, followed by effector T cells. IAV infection led to a classical T cell memory response in the lung, but also increases in IFN*γ*+ ILC1, NK and NKT cells in the draining lymph node, suggesting memory-like phenotypes. These novel findings identify different sources and localisation of IFN*γ* following viral infection which may impact on subsequent infections and improve our understanding of the generation of both classical and non-classical immune memory.

## Materials and Methods

### Study design

The aim of this study was to understand how IFN*γ* production by innate and adaptive immune cells changes over time and following a challenge re-infection viral infection. We used an influenza virus infection model in wildtype and reporter mice and tracked responding memory T cells using MHCI and MHC II tetramers and the reporter systems in the GREATxSMART transgenic mice. A description of the experimental parameters, samples sizes, any samples that were excluded, and the statistical analysis are described in each figure legend. No specified randomisation was conducted.

### Animals

10 week old female C57BL/6 mice were purchased from Envigo (UK). C57BL/6 and GREATxSMART mice, original made by Richard Locksley^29, 47^ and initially provided by David Withers, University of Birmingham, were maintained at the University of Glasgow under specific pathogen free conditions in accordance with UK home office regulations (Project Licenses P2F28B003 and PP1902420) and approved by the local ethics committee. GREATxSMART mice have been described previously^47^.

### Infections

IAV was prepared and titratred in MDCK cells. Mice were briefly anesthetised using inhaled isoflurane and infected with 200 plaque forming units of IAV strain WSN in 20μl of PBS intranasally (i.n.). Infected mice were rechallenged with 200 PFU of X31 (H3N2) where stated. Infected mice were weighed daily for 14 days post-infection. Any animals that lost more than 20% of their starting weight were humanely euthanised.

### Tissue preparation

Mice were injected intravenously (i.v.) with 1μg anti-CD45-PE (ThermoFisher: clone: 30F11) 3 minutes before being euthanized by cervical dislocation. Spleen and mediastinal lymph nodes were processed by mechanical disruption. Single cell suspensions of lungs were prepared by digestion with 1mg/ml collagenase and 30μg/ml DNAse (Sigma) for 40 minutes at 37°C in a shaking incubator. Red blood cells were lysed from spleen and lungs using lysis buffer (ThermoFisher).

### Flow cytometry staining

Single cell suspensions were stained with PE or APC-labelled IA^b^/NP_311-325_ or APC labelled D^b^/ NP_368-374_ tetramers (NIH tetramer core) at 37°C, 5% CO_2_ for 2 hours in complete RPMI (RPMI with 10% foetal calf serum, 100μg/ml penicillin-streptomycin and 2mM L-glutamine) containing Fc block (24G2). Surface antibodies were added and the cells incubated for a further 20 minutes at 4°C. Antibodies used were: anti-CD3 BV785 (BioLegend; clone: 17A2), anti-CD4 BUV805 (BD Bioscience; clone: RM4-5), anti-CD8 BV421 (ThermoFisher; clone: 53-6.7), anti-CD44 BUV395 (BD; clone: IM7), anti-CD45.2 BV605 (BioLegend; clone: 104), anti-CD69 PE-Cy7 (ThermoFisher; clone: H1.2F3) anti-CD127 APC (ThermoFisher; clone: A7R34), anti-*γδ*TCR PE-Cy7 (BioLegend; clone: GL3), anti-ICOS BV785 (BioLegend; clone: C398.4A), anti-NK1.1 APC-Cy7 (BioLegend; clone: PK136), anti-PD1 BV711 (BioLegend; clone: 29F,1A12), and ‘dump’ antibodies: B220 (RA3-6B2), F4/80 (BM8) and MHC II (M5114) all on eFluor-450 (ThermoFisher) or PerCP-Cy5.5 (ThermoFisher; B220 and F4/80, and BioLegend; MHCII). Cells were stained with a fixable viability dye eFluor 506 (ThermoFisher). Stained cells were acquired on a BD LSR or Fortessa and analysed using FlowJo (version 10, Treestar).

### FACS

T cells were isolated from single cell suspensions of mediastinal lymph nodes and spleens using a T cell isolation kit as per manufacturer’s instructions (Stem Cell). Cells were stained with surface antibodies and sorted on a FACS Aria IIU. CD4+ or CD8+ TCRβ+ CD44hi cells that were EYFP+ or EYFP negative were sorted into Qiagen RLTbuffer and stored at −80°C.

### qPCR

RNA was extracted following the manufacturer’s instructions (Qiagen RNAeasy microkit) and analysed by qPCR (SYBR Green FasstMix (Quanta Bioscience). *Ifn*γ** and *18s* standards were generated from spleen cells from IAV infected mice (Primers: *Ifn*γ**: Forward: ATCTGGAGGAACTGGCAAAA; Reverse: AGATACAACCCCGCAATCAC; *18s* Forward: CGTAGTTCCGACCATAAACGA; Reverse: ACATCTAAGGGCATCACAGACC) and purified by gel extraction (Quick Gel Extraction kit, Invitrogen). qPCR was performed on a QuantStudio 7 flex and expression calculated using standard curves and results normalised to *18s* expression. qPCR Primers: *Ifn*γ**: Forward: AGCAAGGCGAAAAAGGATG; Reverse: CTGGACCTGTGGGTTGTTG; *18s*: Forward: GACTCAACACGGGAAACCTC; Reverse: TAACCAGACAAATCGCTCCAC.

### Lung immunofluorescence imaging

Lungs from GREATxSMART mice were perfused with 5mM EDTA and 1% PFA prior to removal and incubated at 4°C in 1% PFA and 30% sucrose for 24 hours each, frozen in OCT and stored at −80°C. 10µm lung sections were cut onto SuperFrost microscope slides (ThermoFisher) and stored at −20°C prior to staining. Sections were incubated in Fc block (24G2) for 10 minutes to block non-specific binding, washed in 0.5% BSA/PBS (ThermoFisher) and stained with antibodies overnight. Antibodies used were: anti-GFP (Life Technologises), anti-CD4 AF647 (BD; clone: RM4-5), anti-CD8b APC (BioLegend, clone; 53-5.8), anti-EpCAM AF594 (BioLegend: G8.8) and anti-MHCII eFluor450 (ThermoFisher; clone: M5114), slides were washed in 0.5% BSA/PBS and mounted using Vectashield mounting reagent (Vector Laboratories). Immunofluorescence images were acquired using a Zeiss LSM800 microscope, analysed using Volocity (version 7, Quorum Technologies) and example images generated on Zen 2 lite.

### Statistical analysis

Data were analysed using Prism version 9 software (GraphPad). Groups were tested for normality using a Shapiro Wilk test, and differences between groups were analysed by one-way ANOVAs, Kruskall-Wallis or unpaired t-tests as indicated in figure legends. In all figures based on distribution normality, * represents a p value of <0.05; **: p>0.01, ***: p>0.001, ****: p>0.0001.

### CRediT authorship contribution statement

GEF: conception, investigation, formal analysis, visualization, writing -original draft and reviewing/editing. KEH: investigation, data curation, writing – reviewing and editing. MP: investigation, data curation, formal analysis, writing – reviewing and editing. TP: investigation, writing – reviewing and editing. DT: formal analysis, writing – reviewing and editing. FM: formal analysis, writing – reviewing and editing. JCW: Supervision, investigation, formal analysis, writing – reviewing and editing. MKLM: Supervision, project management, funding acquisition, conception, investigation, formal analysis, visualization, writing -original draft and reviewing.

## Acknowledgments

We thank the staff within the School of Infection and Immunity Flow Cytometry Facility and Biological Services at the University of Glasgow for technical assistance. We thank David Withers for GREATXSMART mice and the NIH tetramer core facility for the provision of IA^b^-NP_311-325_ and Db/NP NP_368-74_ tetramers. The work was funded by a University of Glasgow MVLS PhD studentship awarded to GEF and a Wellcome Trust Investigator Award (210703/Z/18/Z) to MKLM.

---

**Supplementary Figure 1.**
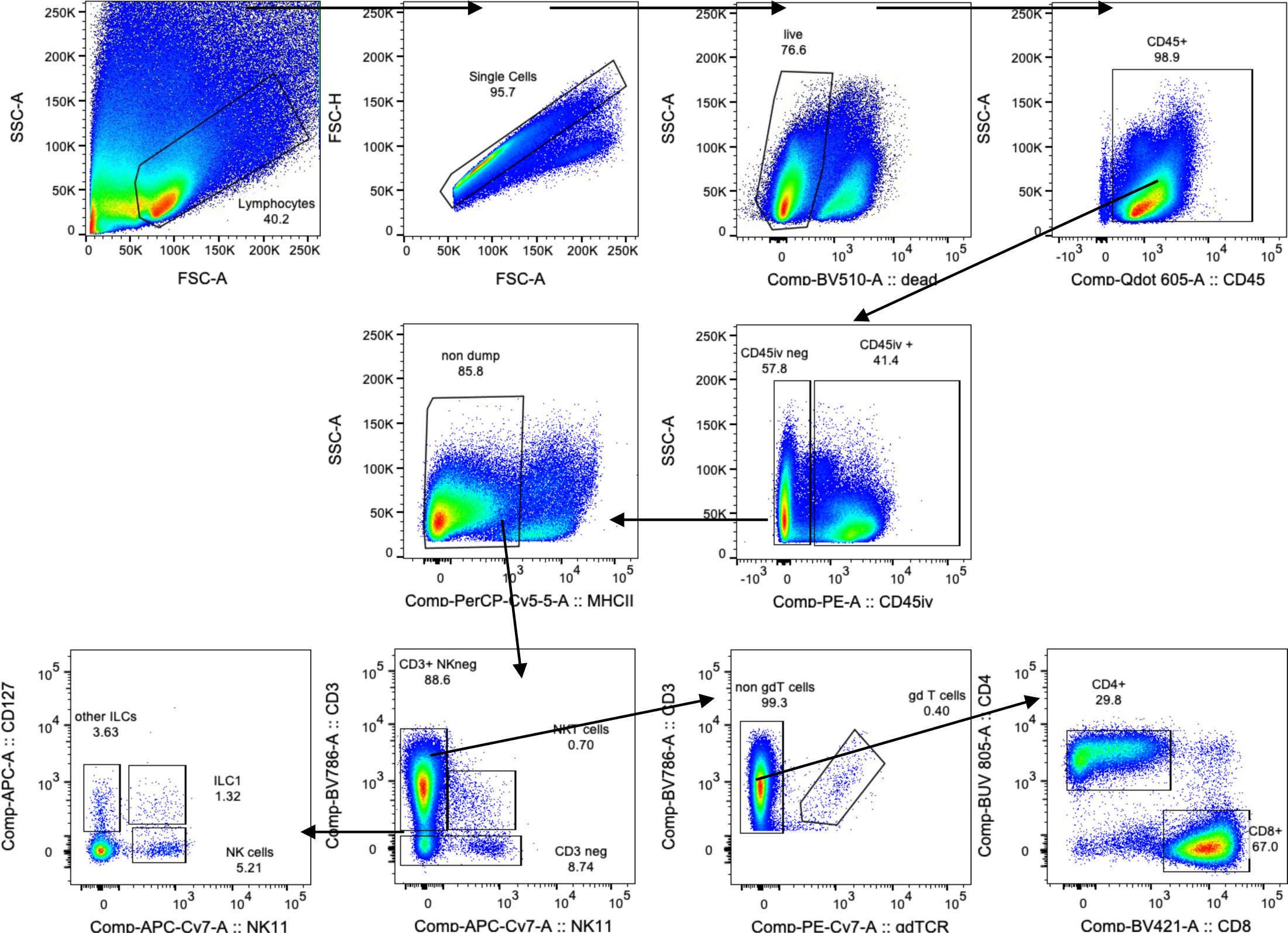
Gating strategy diagram for flow cytometry analysis of immune cell populations in IAV-infected mouse lung. GREATxSMART mice were infected with IAV on day 0 and injected with fluorescently labelled anti-CD45 i.v. 3 minutes prior to removal of the tissues. Representative flow for NK cells (CD3-, NK1.1+, CD127-), NK T cells (CD3+, NK1.1+), ILC1s (CD3-, NK1.1+, CD127) other ILCs (CD3-, NK1.1-, CD127+), CD4 (CD3+, ***γδ*** TCR-, CD4+, CD8-) CD8 (CD3+, ***γδ*** TCR-, CD4-, CD8+) and ***γδ*** T cells (CD3+, ***γδ*** TCR+) 10 days following infection in the lung. Numbers indicate percentage of IFN*γ*^+^ populations of each cell type.

**Supplementary Figure 2.**
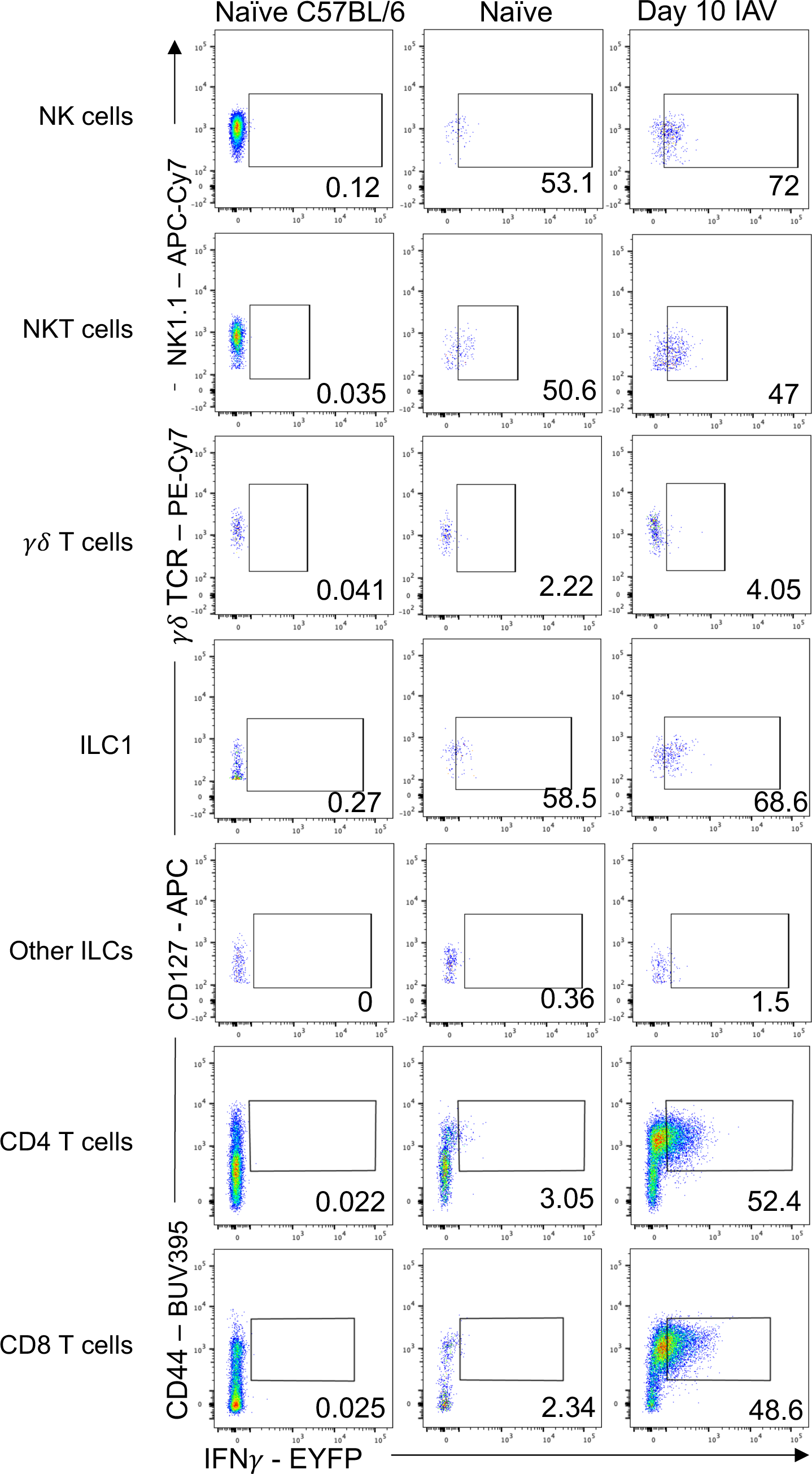
Validation of GREATxSMART mouse EYFP expression. GREATxSMART mice were infected with IAV on day 0 and injected with fluorescently labelled anti-CD45 i.v. 3 minutes prior to removal of tissues. Single cell suspensions of lungs were examined after 10 days to examine IFN*γ* transcription of indicated cell populations determined by EYFP+ CD45iv^-^ populations compared to naïve and naïve C57BL/6 mice. Numbers indicate percentage of IFN*γ*^+^ populations of each cell type.

**Supplementary Figure 3.**
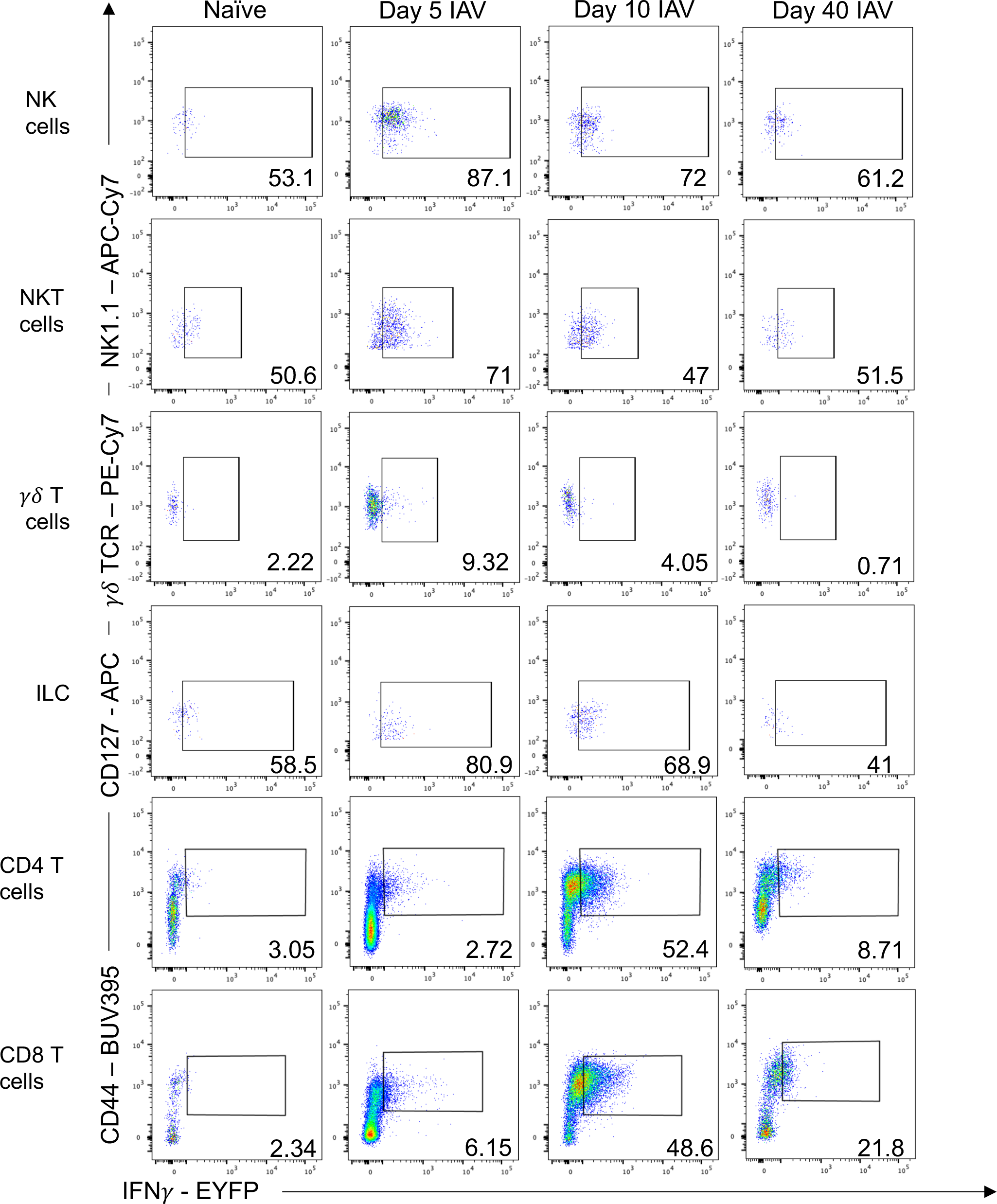
Characterisation of IFN*γ* expression from immune cells during IAV infection in the lung. GREATxSMART mice were infected with IAV on day 0 and injected with fluorescently labelled anti-CD45 i.v. 3 minutes prior to removal of the tissues. Single cell suspensions of lungs were examined after 5, 10 or 40 days to examine IFN*γ* transcription of indicated cell populations determined by EYFP+ and CD45iv^-^ populations compared to naïve. Numbers indicate percentage of IFN*γ*^+^ populations of each cell type.

**Supplementary Figure 4.**
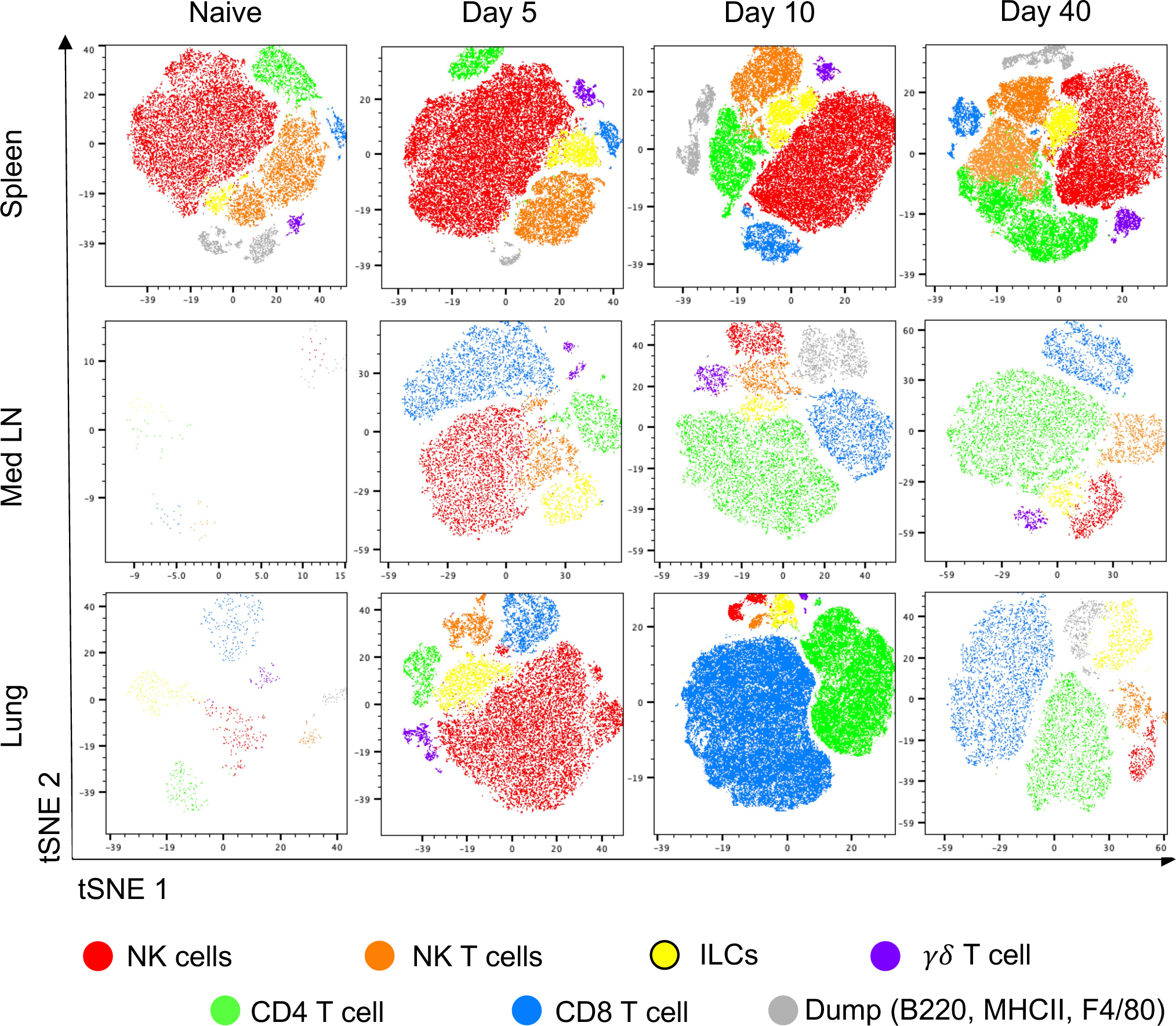
Validation of GREATxSMART mouse ability to detect IFN*γ* expression through EYFP fluorescence from various immune cells. Representative tSNE plots depicting IFN*γ*+ clusters identified from GREATxSMART mice that were infected with IAV and 5, 10 or 40 days later. Mice were injected with fluorescently labelled anti-CD45 shortly before tissues were harvested. IFN*γ*-producing cells within the spleen, Med LN, and lung were examined by flow cytometry and determined by EYFP+ and CD45iv^-^ populations compared to naïve. Cell types identified are NK cells (red), NKT cells (orange), ILCs, (yellow), CD4 T cells (green), CD8 T cells (blue), ***γδ*** T cells (purple) and dump (B220, MHCII and F4/80) channel cells (grey).

**Supplementary Figure 5.**
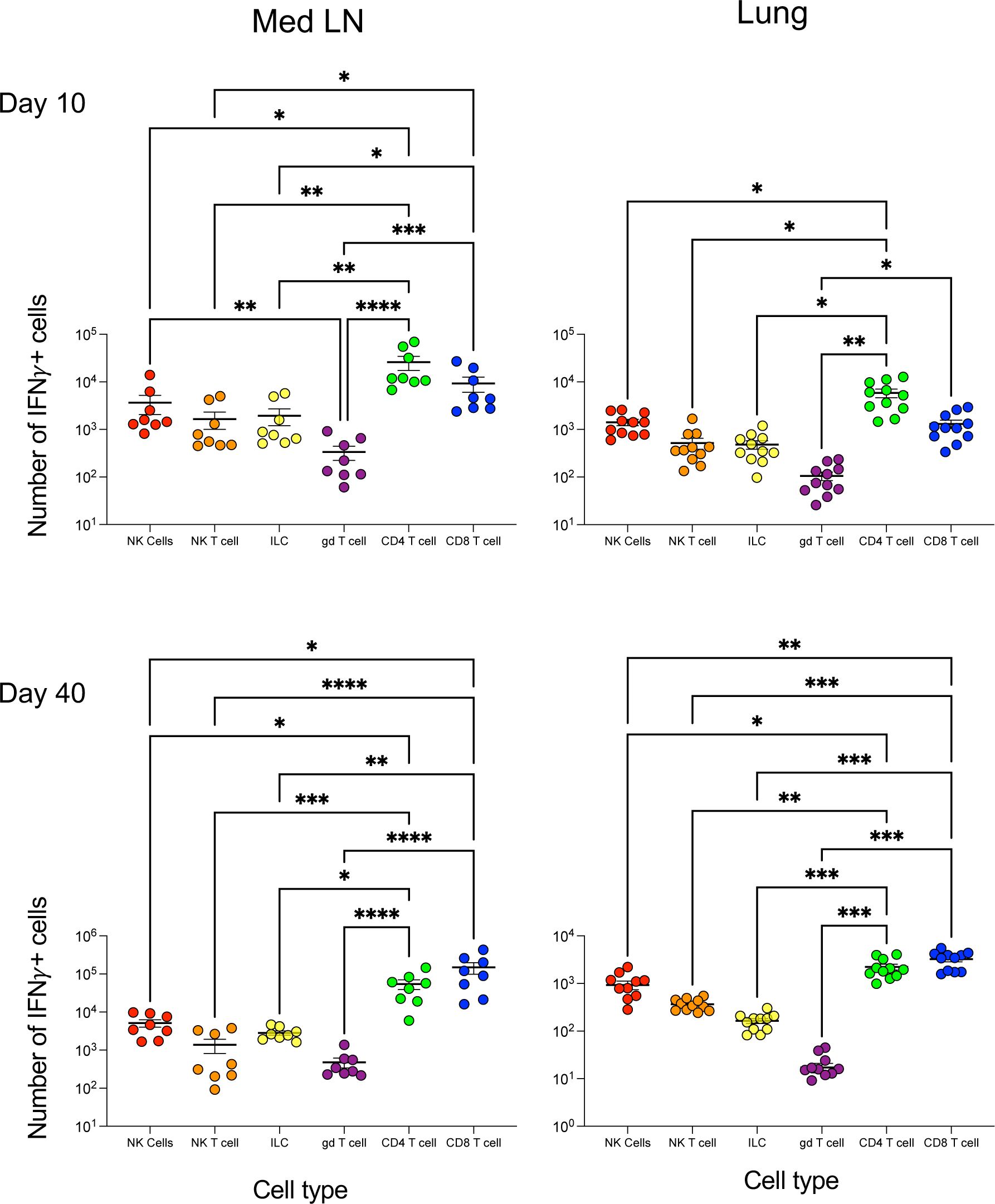
CD4 and CD8 T cells are the predominant source of IFN*γ* 10 and 40 days after IAV infection in the lung and lymph node. GREATxSMART mice were infected with IAV on day 0 and injected with fluorescently labelled anti-CD45 i.v. 3 minutes prior to removal of the tissues. The total number of IFN*γ*+ cells of the indicated cell populations in the Med LN and lung were examined in animals infected 10 (top) or 40 (bottom) days previously. Each point represents an individual mouse of 8-11 infected mice from two independent time course experiments; 7-11 mice are combined from across the time points and experiments, error bars are SEM. All statistics were calculated using a Shapiro-Wilk and Kruskal-Wallis test., * = <0.05, ** = <0.01, *** = <0.001, **** = <0.0001.

**Supplementary Figure 6.**
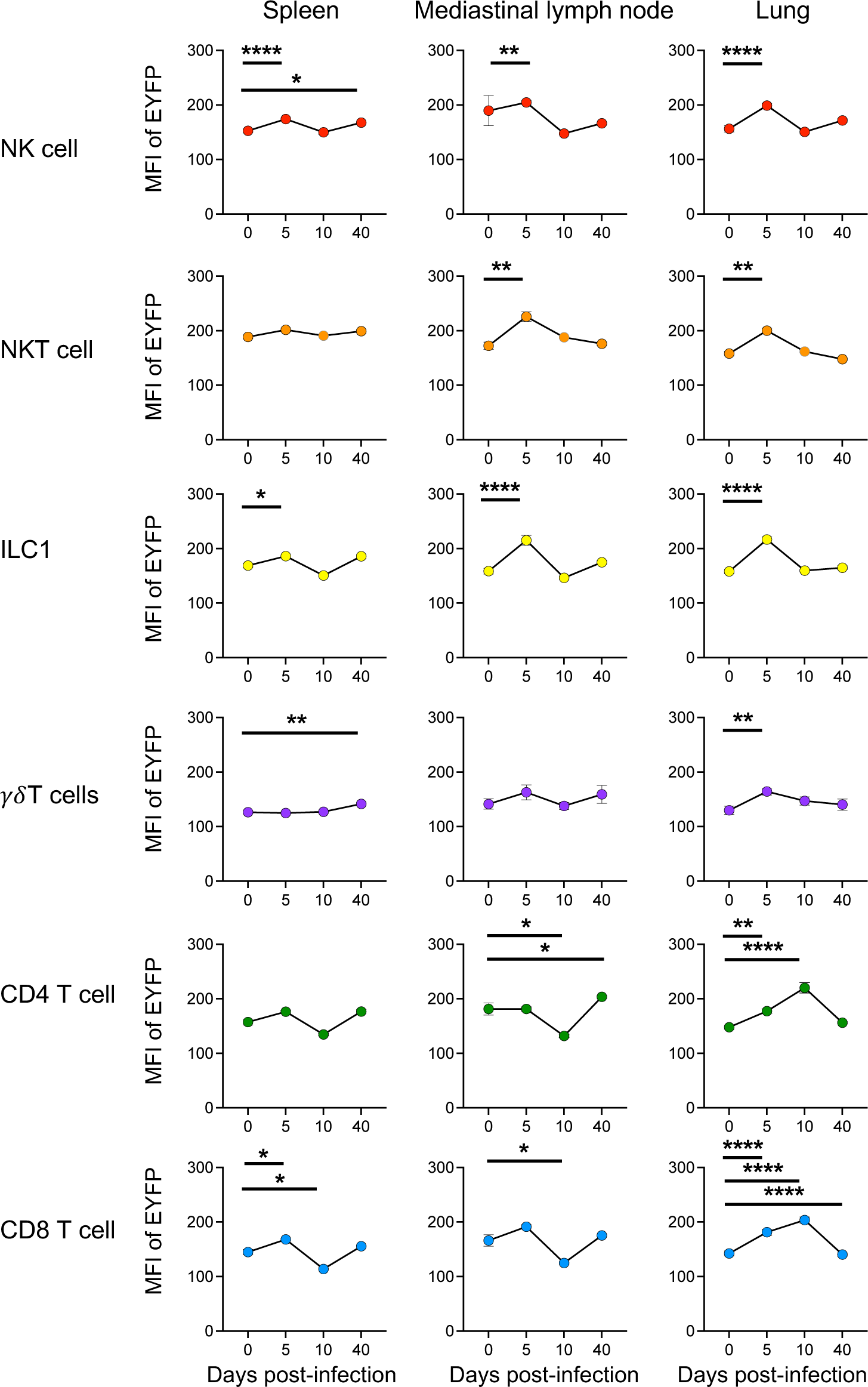
The amount of immune cell-derived IFN*γ* fluctuates between the tissue and secondary lymphoid organs. GREATxSMART mice were infected with IAV on day 0 and injected with fluorescently labelled anti-CD45 i.v. 3 minutes prior to removal of the spleen, Med LN and lung. The mean fluorescence intensity (MFI) of the indicated cell populations were examined in naïve animals or those infected 5, 10 or 40 days previously. Each point represents the mean of 8-11 infected mice from two independent time course experiments; 21 naïve mice are combined from across the time points and experiments, error bars are SEM. Significance tested via a Kruskal–Wallis test followed by a Dunn’s multiple comparison test, *: p<0.05.

**Supplementary Figure 7.**
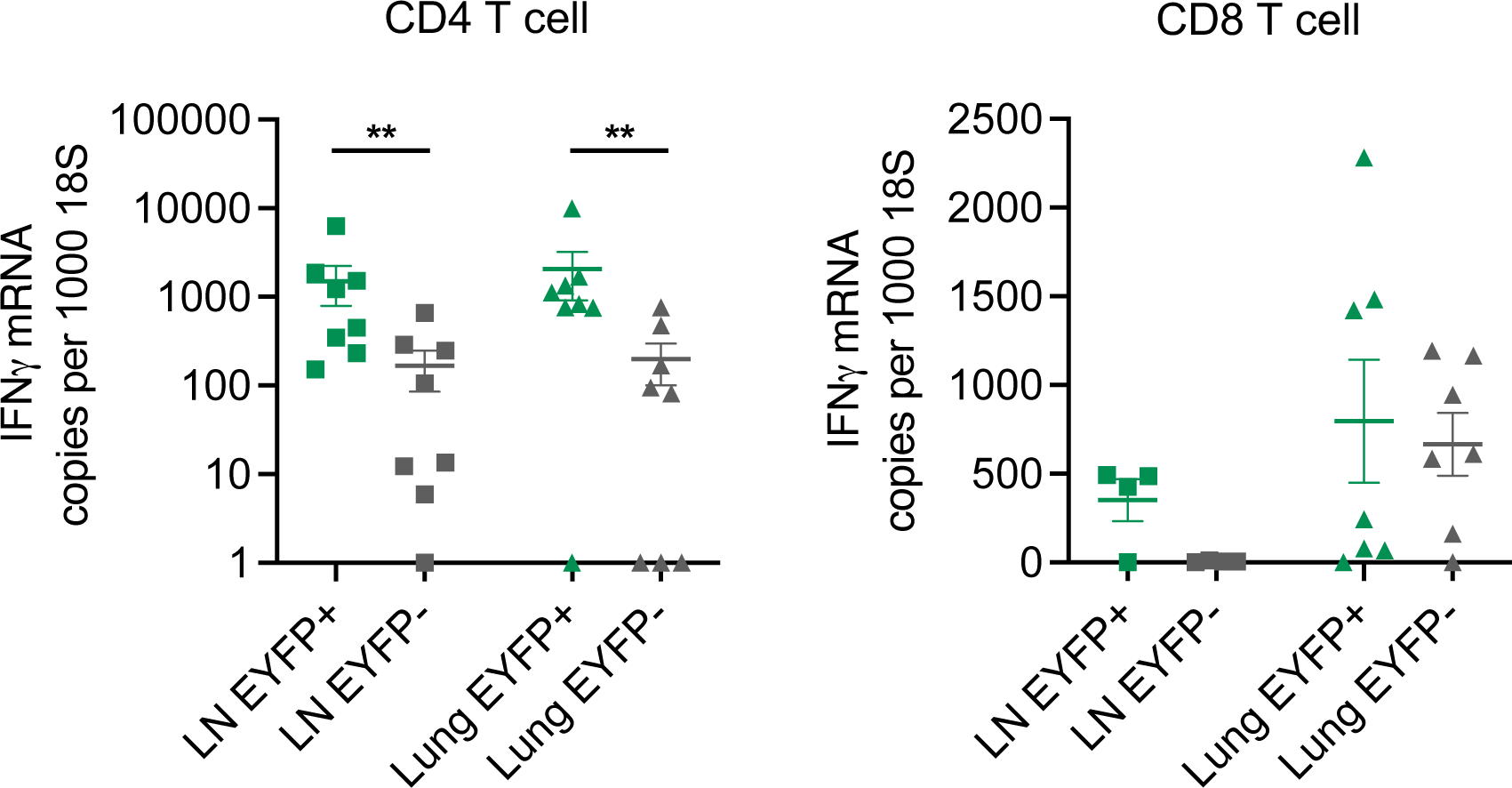
EYFP expression reports IFN*γ* expression in CD4 T cells after IAV infection. GREATxSMART mice were infected with 200PFU IAV i.n and EYFP+ and EYFP-T cells were FACS sorted from the Med LN and lung of mice 40 days after infection. Gene expression of IFN*γ* by CD4 (left) and CD8 (right) T cells in the lymph node (LN) and lung of IAV-infected mice shown. All gene expression was standardised against the housekeeping gene (18S) and is presented as the change in absolute copy number of transcripts. Each point represents an individual mouse and data are combined from two independent experiments; error bars are SEM. Significance tested via a Shapiro-Wilk normality test and Wilcoxon test. ** = <0.01.

**Supplementary Table 1.**
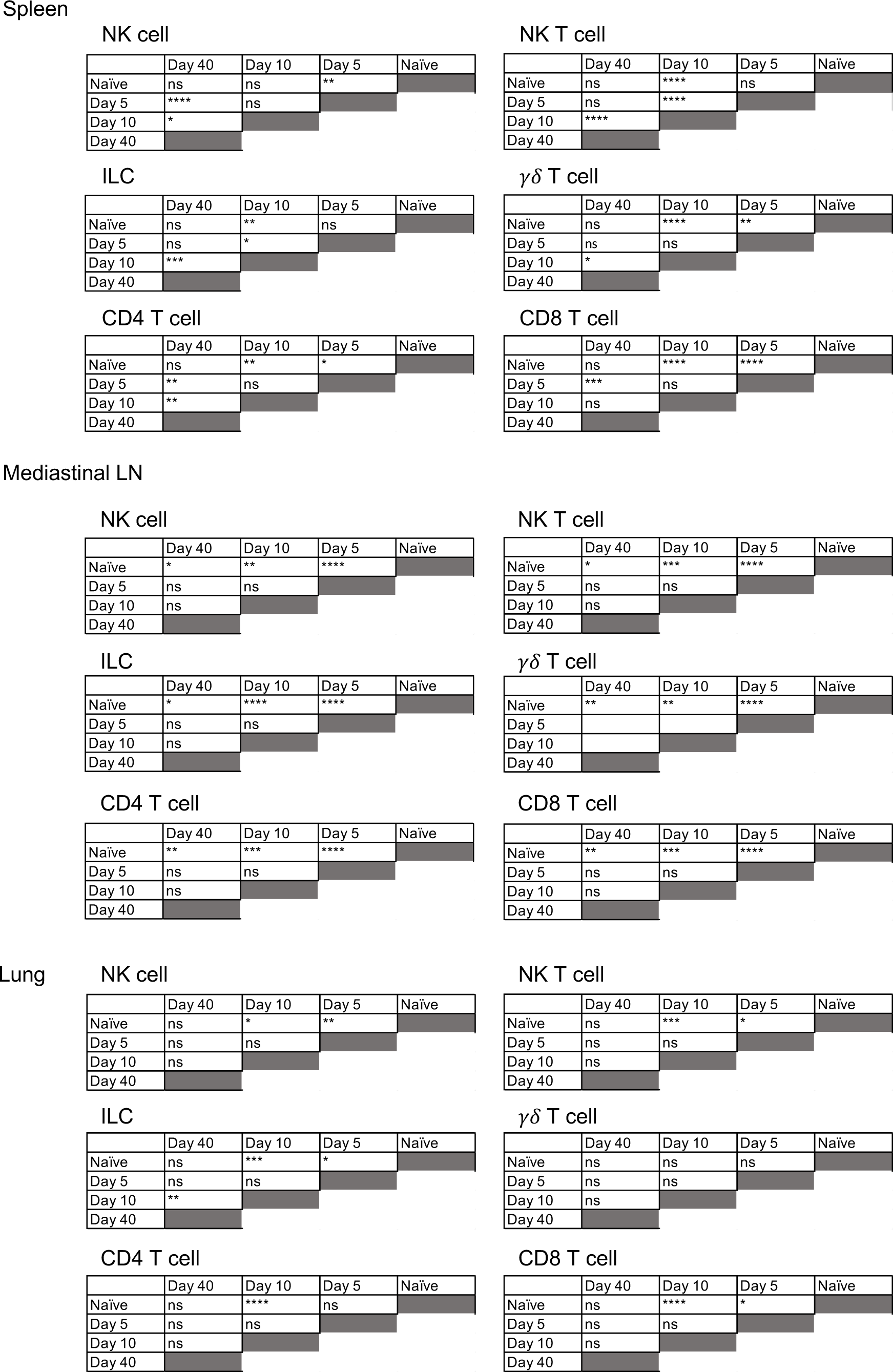
Characterization of immune cells during IAV infection. Significance tables from spleen, draining lymph node and lungs from mice infected with IAV, comparing significantly different numbers of NK cells, NK T cells, ILCs, *γδ* T cells, CD4 and CD8 T cells. Significance tested via a Kruskal–Wallis test followed by a Dunn’s multiple comparison test, ns = non-significant, *: p<0.05, **: p<0.01, ***: p<0.001, ****: p<0.0001.

**Supplementary Table 2.**
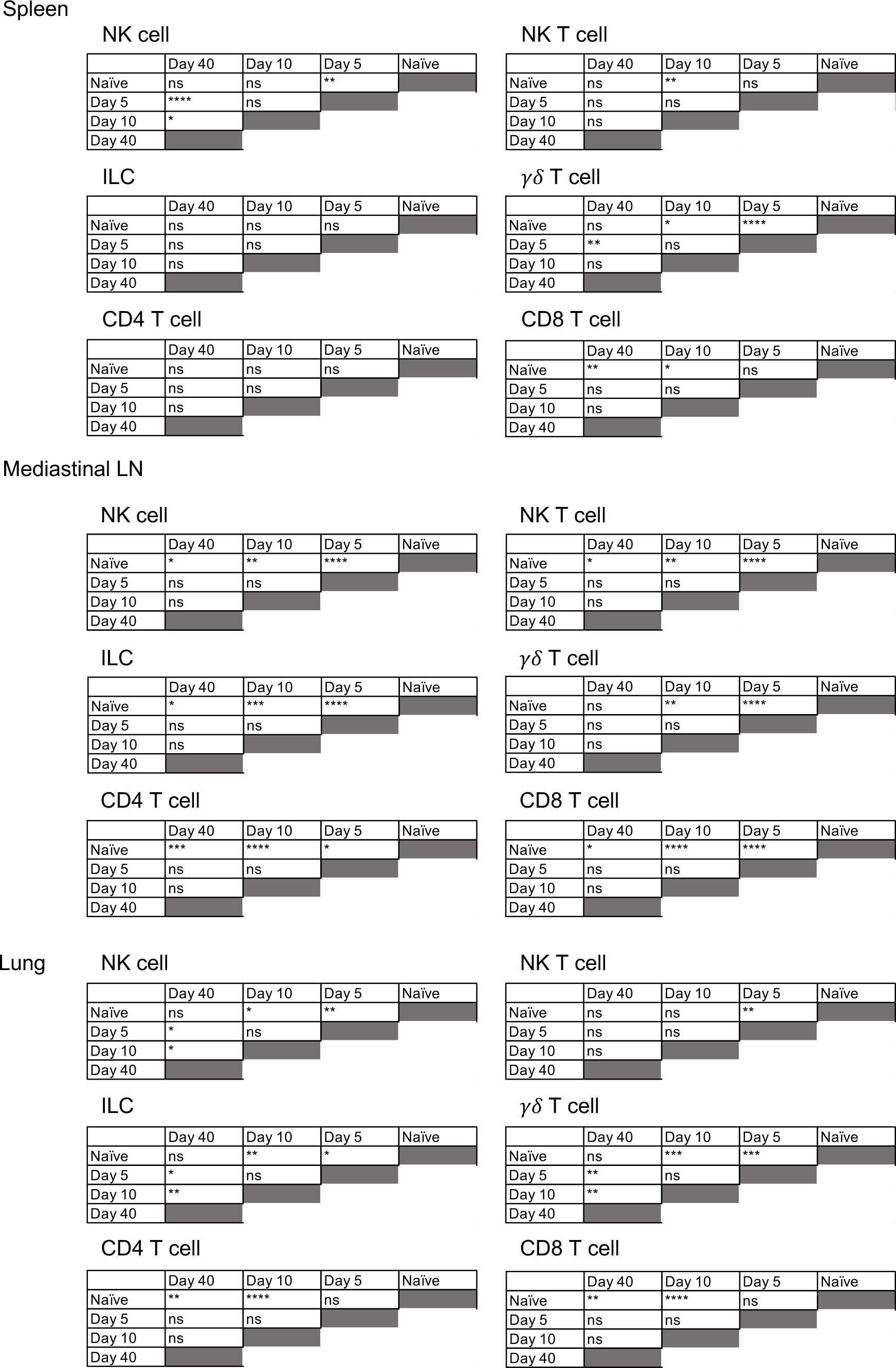
Characterization of IFN*γ*-expressing cells during IAV infection. Significance tables from spleen, draining lymph node and lungs from mice infected with IAV, comparing significantly different numbers of IFN*γ*+ NK cells, NK T cells, ILCs, *γδ* T cells, CD4 and CD8 T cells. Significance tested via a Kruskal–Wallis test followed by a Dunn’s multiple comparison test, ns = non-significant, * = <0.05, ** = <0.01, *** = <0.001, **** = <0.0001.

**Supplementary Table 3.** Characterization of the mean fluorescence intensity of IFN*γ* produced by cells during IAV infection. Significance tables from spleen, draining lymph node and lungs from mice infected with IAV, comparing significantly different mean fluorescence intensity (MFI) of IFN*γ*+ NK cells, NK T cells, ILCs, *γδ* T cells, CD4 and CD8 T cells. Significance tested via a Kruskal–Wallis test followed by a Dunn’s multiple comparison test, ns = non-significant, * = <0.05, ** = <0.01, *** = <0.001, **** = <0.0001.

|  | Day 40 | Day 10 | Day 5 | Naïve |
| --- | --- | --- | --- | --- |
| Naïve | * | ns | **** |  |
| Day 5 | ns | **** |  |  |
| Day 10 | * |  |  |  |
| Day 40 |  |  |  |  |

ILC
|  | Day 40 | Day 10 | Day 5 | Naïve |
| --- | --- | --- | --- | --- |
| Naïve | ns | ns | * |  |
| Day 5 | ns | *** |  |  |
| Day 10 | *** |  |  |  |
| Day 40 |  |  |  |  |

CD4 T cell
|  | Day 40 | Day 10 | Day 5 | Naïve |
| --- | --- | --- | --- | --- |
| Naïve | ns | ns | ns |  |
| Day 5 | ns | ** |  |  |
| Day 10 | ** |  |  |  |
| Day 40 |  |  |  |  |

$\gamma\delta$ T cell
|  | Day 40 | Day 10 | Day 5 | Naïve |
| --- | --- | --- | --- | --- |
| Naïve | ** | ns | ns |  |
| Day 5 | ** | ns |  |  |
| Day 10 | * |  |  |  |
| Day 40 |  |  |  |  |

CD8 T cell
|  | Day 40 | Day 10 | Day 5 | Naïve |
| --- | --- | --- | --- | --- |
| Naïve | ns | * | * |  |
| Day 5 | ns | **** |  |  |
| Day 10 | * |  |  |  |
| Day 40 |  |  |  |  |

Mediastinal LN NK cell
|  | Day 40 | Day 10 | Day 5 | Naïve |
| --- | --- | --- | --- | --- |
| Naïve | ns | ns | ** |  |
| Day 5 | ns | ns |  |  |
| Day 10 | **** |  |  |  |
| Day 40 |  |  |  |  |

ILC
|  | Day 40 | Day 10 | Day 5 | Naïve |
| --- | --- | --- | --- | --- |
| Naïve | ns | ns | **** |  |
| Day 5 | **** | **** |  |  |
| Day 10 | ** |  |  |  |
| Day 40 |  |  |  |  |

CD4 T cell
|  | Day 40 | Day 10 | Day 5 | Naïve |
| --- | --- | --- | --- | --- |
| Naïve | * | * | ns |  |
| Day 5 | ns | ns |  |  |
| Day 10 | **** |  |  |  |
| Day 40 |  |  |  |  |

NK T cell
|  | Day 40 | Day 10 | Day 5 | Naïve |
| --- | --- | --- | --- | --- |
| Naïve | ns | ns | ** |  |
| Day 5 | ** | ns |  |  |
| Day 10 | ns |  |  |  |
| Day 40 |  |  |  |  |

CD8 T cell
|  | Day 40 | Day 10 | Day 5 | Naïve |
| --- | --- | --- | --- | --- |
| Naïve | ns | * | ns |  |
| Day 5 | ns | *** |  |  |
| Day 10 | ns |  |  |  |
| Day 40 |  |  |  |  |

Lung NK cell
|  | Day 40 | Day 10 | Day 5 | Naïve |
| --- | --- | --- | --- | --- |
| Naïve | ns | ns | **** |  |
| Day 5 | ns | *** |  |  |
| Day 10 | ns |  |  |  |
| Day 40 |  |  |  |  |

ILC
|  | Day 40 | Day 10 | Day 5 | Naïve |
| --- | --- | --- | --- | --- |
| Naïve | ns | ns | **** |  |
| Day 5 | ** | ** |  |  |
| Day 10 | ns |  |  |  |
| Day 40 |  |  |  |  |

CD4 T cell
|  | Day 40 | Day 10 | Day 5 | Naïve |
| --- | --- | --- | --- | --- |
| Naïve | ns | **** | ** |  |
| Day 5 | ns | ns |  |  |
| Day 10 | ** |  |  |  |
| Day 40 |  |  |  |  |

NK T cell
|  | Day 40 | Day 10 | Day 5 | Naïve |
| --- | --- | --- | --- | --- |
| Naïve | ns | ns | ** |  |
| Day 5 | **** | ns |  |  |
| Day 10 | ns |  |  |  |
| Day 40 |  |  |  |  |

|  | Day 40 | Day 10 | Day 5 | Naïve |
| --- | --- | --- | --- | --- |
| Naïve | *** | **** | **** |  |
| Day 5 | **** | ns |  |  |
| Day 10 | *** |  |  |  |
| Day 40 |  |  |  |  |

